# Potent mouse monoclonal antibodies that block SARS-CoV-2 infection

**DOI:** 10.1101/2020.10.01.323220

**Authors:** Youjia Guo, Atsushi Kawaguchi, Masaru Takeshita, Takeshi Sekiya, Mikako Hirohama, Akio Yamashita, the Keio Donner Project, Haruhiko Siomi, Kensaku Murano

## Abstract

Coronavirus disease 2019 (COVID-19), caused by severe acute respiratory syndrome coronavirus 2 (SARS-CoV-2), has developed into a global pandemic since its first outbreak in the winter of 2019. An extensive investigation of SARS-CoV-2 is critical for disease control. Various recombinant monoclonal antibodies of human origin that neutralize SARS-CoV-2 infection have been isolated from convalescent patients and will be applied as therapies and prophylaxis. However, the need for dedicated monoclonal antibodies in molecular pathology research is not fully addressed. Here, we produced mouse anti-SARS-CoV-2 spike monoclonal antibodies that exhibit not only robust performance in immunoassays including western blotting, ELISA, immunofluorescence, and immunoprecipitation, but also neutralizing activity against SARS-CoV-2 infection *in vitro*. Our monoclonal antibodies are of mouse origin, making them compatible with the experimental immunoassay setups commonly used in basic molecular biology research laboratories, and large-scale production and easy distribution are guaranteed by conventional mouse hybridoma technology.

## Introduction

The outbreak of COVID-19 caused by severe acute respiratory syndrome coronavirus 2 (SARS-CoV-2) is a threat to global public health and economic development (Huang et al., 2020; Li et al., 2020). Vaccine and therapeutic discovery efforts are paramount to restrict the spread of the virus. Passive immunization could have a major effect on controlling the virus pandemic by providing immediate protection, complementing the development of prophylactic vaccines (Klasse & Moore, 2020; Walker & Burton, 2018). Passive immunization against infectious diseases can be traced back to the late 19th century and the work of Shibasaburo Kitasato and Emil von Behring on the serotherapy of tetanus and diphtheria. There have been significant developments in therapies and prophylaxis using antibodies over the past 50 years (Graham & Ambrosino, 2015).

The advent of hybridoma technology in 1975 provided a reliable source of mouse monoclonal antibodies (Kohler & Milstein, 1975). With the development of humanized mouse antibodies and subsequent generation of fully human antibodies by various techniques, monoclonal antibodies have become widely used in therapy and prophylaxis for cancer, autoimmune diseases, and viral pathogens (Walker & Burton, 2018). Indeed, a humanized mouse monoclonal antibody neutralizing respiratory syncytial virus (RSV), palivizumab, is widely used in clinical settings prophylactically to protect vulnerable infants (Connor, 1999). In recent years, highly specific and often broadly active neutralizing monoclonal antibodies have been developed against several viruses (Caskey, Klein, & Nussenzweig, 2019; Corti et al., 2017; Davide Corti et al., 2016; Corti, Passini, Lanzavecchia, & Zambon, 2016; Walker & Burton, 2018). Passive immunization with a monoclonal antibody is currently under consideration as a treatment for COVID-19 caused by SARS-CoV-2 (Dhama et al., 2020; Jawhara, 2020; Jiang, Hillyer, & Du, 2020; Klasse & Moore, 2020; Ni et al., 2020).

Isolation of multiple human neutralizing monoclonal antibodies against SARS-CoV-2 has been reported (Cao et al., 2020; Chen et al., 2020; Chi et al., 2020; Hassan et al., 2020; Ju et al., 2020; Liu et al., 2020; Pinto et al., 2020; Robbiani et al., 2020; Rogers et al., 2020; Shi et al., 2020; Wan et al., 2020; Wang et al., 2020; Wu et al., 2020; Zeng et al., 2020; Zost et al., 2020). These antibodies can avoid the potential risks of human-anti-mouse antibody responses and other side effects (Hansel, Kropshofer, Singer, Mitchell, & George, 2010). They will be appropriate for direct use in humans since they are humanized even if these monoclonal antibodies are recombinant. Owing to the recent rapid development of single-cell cloning technology, the process of antibody isolation has been dramatically shortened compared with the generation of a conventional monoclonal antibody secreted from a hybridoma resulting from the fusion of a mouse myeloma with B cells (Wan et al., 2020). However, since they are recombinant human antibodies produced in HEK293 cell lines derived from human embryonic kidney, they have a disadvantage compared to conventional hybridoma-produced antibodies in terms of their lot-to-lot quality control and manufacturing costs (Cohen, 2020). Instead, monoclonal antibodies produced by hybridomas are secreted into the culture supernatant, thus their production is straightforward and of low cost, and their quality is stable. It is also easy to distribute them to researchers worldwide, although they will not be applicable for treatment, if not chimeric and humanized, due to their immunogenicity (Hansel et al., 2010; Reichert, Rosensweig, Faden, & Dewitz, 2005).

In addition to the impact of monoclonal antibodies on therapy and prophylaxis, they significantly impact the characterization of SARS-CoV-2. To overcome the long-term battle with the virus, we need a detailed understanding of the replication mechanisms underlying its lifecycle, including viral entry, genome replication, budding from the cellular membrane, and interaction with host immune systems. These essential pieces of information are required for drug discovery, vaccine design, and therapy development. Despite the large number of neutralizing antibodies reported to inhibit infection, there is an overwhelming lack of data on a well-characterized antibody available for basic research techniques such as western blotting, immunofluorescence, and immunoprecipitation to study the viral life cycle.

Here, we established six monoclonal antibodies against the spike glycoprotein of SARS-CoV-2. The trimeric spike glycoproteins of SARS-CoV-2 play a pivotal role in viral entry into human target cells through the same receptor, angiotensin-converting enzyme 2 (ACE2) as SARS-CoV-1 (Hoffmann et al., 2020). Our antibodies were produced by a hybridoma resulting from the fusion of a mouse myeloma SP2/0 with splenocytes obtained from BALB/c mice immunized with purified recombinant spike proteins. We evaluated these antibodies for application in molecular pathology research. Among them, two antibodies were shown to attenuate the interaction of spike proteins with ACE2 and neutralized infection of VeroE6/TMPRSS2 cells by SARS-CoV-2. Our antibodies will accelerate research on SARS-CoV-2 and lead to new therapies and prophylaxis.

## Results

### Production of six monoclonal antibodies against spike glycoprotein

The SARS-CoV-2 spike glycoprotein is a homotrimeric fusion protein composed of two subunits: S1 and S2. During infection, the receptor binding domain (RBD) on S1 subunit binds to ACE2, resulting in destabilization of the spike protein’s metastable conformation. Once destabilized, the spike protein is cleaved into the N-terminal S1 and C-terminal S2 subunits by host proteases such as TMPRSS2 and changes conformation irreversibly from the prefusion to the postfusion state (Hoffmann et al., 2020; Ou et al., 2020; Song, Gui, Wang, & Xiang, 2018), which triggers an infusion process mediated by the S2 region (Tai, Zhang, He, Jiang, & Du, 2020; Walls et al., 2020). The instability needs to be addressed to obtain high-quality spike proteins for downstream applications. We adopted the design principle reported by Wrapp *et al*. (Wrapp et al., 2020), in which the SARS-CoV-2 spike protein was engineered to form a stable homotrimer that was resistant to proteolysis during protein preparation. In our practice, recombinant spike protein RBD and ectodomain were constructed. A T4 fabritin trimerization motif (foldon) was incorporated into the C-terminal of the recombinant spike ectodomain to promote homotrimer formation (Miroshnikov et al., 1998) (Fig. 1A). Recombinant RBD proteins tagged with GST or MBP were produced using an *E. coli* expression system (Fig. 1B). Both recombinant spike protein RBD and ectodomain (SΔTM) were produced using a mammalian expression system that retained proper protein glycosylation equivalent to that observed during virus replication (Fig. 1C, S1A). Mice were immunized with these recombinant spike proteins to generate antibodies against the SARS-CoV-2 virus, followed by cell fusion to generate a hybridoma-producing antibody. Culture supernatants were pre-screened by enzyme-linked immunosorbent assay (ELISA), western blotting (WB), and immunoprecipitation (IP), and six monoclonal hybridomas were isolated and evaluated.

**Figure 1.**
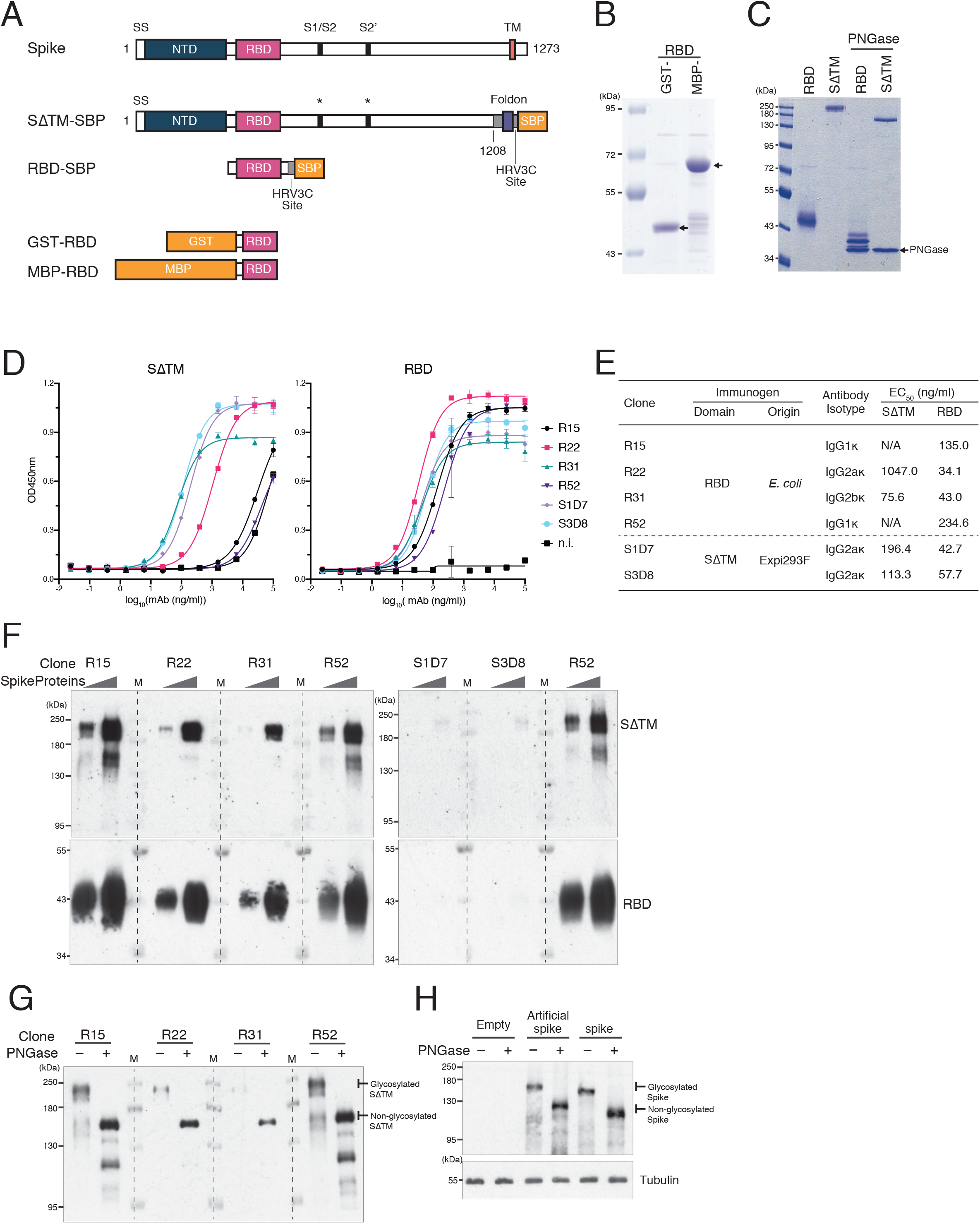
Production of six monoclonal antibodies against spike protein. A. Schematic of recombinant proteins used to establish anti-spike antibodies SS, signal peptide; NTD, N-terminal domain; RBD, receptor-binding domain; TM, transmembrane domain; SΔTM, spike lacking TM domain; SBP, streptavidin binding peptide; Foldon, T4 fabritin trimerization motif; GST, glutathione S-transferase; MBP, maltose binding protein. For mammalian expression constructs (SΔTM-SBP and RBD-SBP), the HRV3C cleavage site was placed upstream of the SBP tag so that the SBP tag could be removed by HRV3C protease treatment after protein purification (Fig. S1A). B. Coomassie brilliant blue (CBB) staining of recombinant protein purified from *E. coli* expression system. GST-RBD and MBP-RBD appeared as bands of 46 kDa and 62 kDa, respectively. C. CBB staining of recombinant proteins purified from the mammalian expression system. The glycosylation of recombinant proteins caused smear bands and a lower migration rate of proteins on SDS-PAGE compared to proteins treated with PNGase. D. ELISA-binding affinity of purified monoclonal antibodies to trimeric SΔTM and RBD glycoproteins purified from the mammalian expression system. n.i., non-immune mouse IgG Error bars indicate standard deviation (n=3). E. Summary of isotype and EC_50_ of established monoclonal antibodies. F. Western blotting (WB) against SΔTM and RBD glycoproteins (10 or 50 ng per lane) using purified monoclonal antibodies (1 µg/mL in PBS-T). Clone S1D7 and S3D7 could not detect either SΔTM or RBD in WB. G. Detection of non-glycosylated SΔTM using established monoclonal antibodies. Four clones could detect SΔTM (30 ng per lane), regardless of glycosylation. H. Detection of spike proteins expressed in 293T cells. Lysates of 293T cells expressing artificial spikes carrying T4 foldon (artificial spike) or wild-type spike glycoproteins were separated by SDS-PAGE, followed by WB using antibody R52.

To characterize these antibodies in detail, they were first purified from the culture supernatant and examined in terms of ELISA and WB performance. Four monoclonal antibodies derived from the antigen produced by *E. coli* (Clones R15, R22, R31, and R52) and two from mammalian cells (S1D7 and S3D8) showed remarkable performance. In the ELISA binding assay, all six clones bound glycosylated RBD with high affinity. When tested against spike glycoprotein (SΔTM), two clones (R15 and R52) could not be distinguished from non-immune IgG (Fig. 1D). We noted that IgG2 subclass members tended to have higher binding affinities. Half maximal effective concentration (EC_50_) required for these antibodies to bind RBD and SΔTM glycoproteins falls at the low hundreds ng/mL (Fig. 1E). In WB, where target proteins are reduced and denatured, all clones established by *E. coli* produced-antigens performed well at detecting RBD and SΔTM proteins regardless of glycosylation (Fig. 1F, left, and 1G, S1B). Among them, clones R15 and R52 showed higher sensitivity in WB. In addition, R52 was capable of detecting not only artificial spike glycoprotein carrying T4 foldon, but also native spike glycoprotein expressed in 293T cells on WB (Fig. 1H). However, neither RBD nor SΔTM could be detected by antibody clones established by the mammalian antigen (S1D7 and S3D8) on WB, suggesting a strong preference for intact tertiary structure (Fig. 1F, right).

### S1D7 and S3D8 antibodies showed higher performance on IP and IF

An antibody capable of recognizing the intact tertiary structure of spike proteins would contribute to research dissecting the molecular mechanism of SARS-CoV-2 infection, especially cell entry, where these proteins play a significant role. The IP activity of antibodies can be correlated with the activity of capturing the native structure of the target protein and neutralizing the infection. We examined the IP performance of our monoclonal antibodies. Although all clones were capable of immunoprecipitating RBD and SΔTM glycoproteins, clone R22, R31, S1D7, and S3D8 demonstrated superior IP efficiency for SΔTM, whereas R22, S1D7, and S3D8 showed higher IP efficiency for RBD glycoprotein (Fig 2A). As shown in Fig. 2B, our antibodies recognize the spike protein in a glycosylation-independent manner, and the IP efficiencies of R22, R31, S1D7, and S3D8, although mild, outperformed others. Noticeably, although clones S1D7 and S3D8 are not capable of performing WB (Fig. 1F), a strong preference for tertiary structure grants them remarkable performance in IP, where RBD and SΔTM glycoproteins were pulled down in their native conformation. Of note, we found that S1D7 and S3D8 could maintain intact IP efficiency under highly stringent experimental conditions where sodium dodecyl sulfate (SDS) was present (Fig S2A).

**Figure 2.**
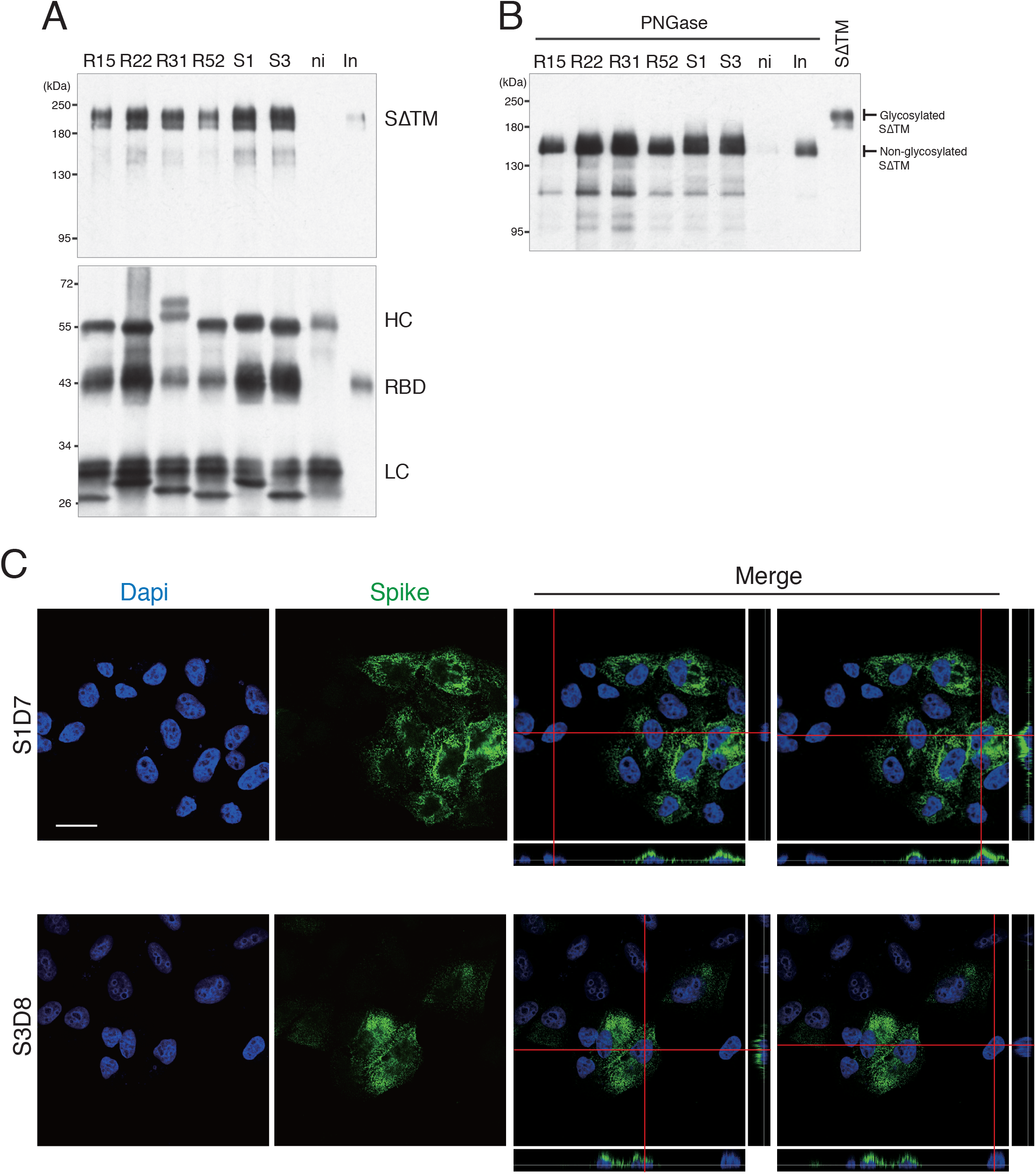
Application for immunoprecipitation and immunofluorescence. A. Immunoprecipitation (IP) of trimeric glycosylated spike protein (SΔTM) using established monoclonal antibodies. S1, S1D7; S3, S3D8; ni, non-immune mouse IgG; In, input; SΔTM, trimeric spike protein without transmembrane domain; HC, IgG heavy chain; LC, IgG light chain. All clones were capable of pulling down RBD and Spike glycoprotein. Higher IP efficiency of Spike glycoprotein was observed in clones R22, R31, S1D7, and S3D8. For RBD glycoprotein, clone R22, S1D7, and S3D8 showed higher IP efficiency. B. IP of trimeric spike protein de-glycosylated by PNGase F using established monoclonal antibodies. “SΔTM” indicates SΔTM glycoprotein untreated with PNGase F. All clones are capable of pulling down de-glycosylated spike protein. Higher IP efficiency was observed in clone R22, R31, S1D7, and S3D8. C. Immunofluorescence (IF) staining of spike glycoprotein expressed in HeLa cells with monoclonal antibodies S1D7 and S3D8. Spike protein localized on the apical surface of transfected HeLa cells Scale bar, 30 µm.

Next, we examined whether our antibodies could be used in the immunofluorescence assay (IF). An antibody applicable for IP would also have activity in IF. Cellular localization of spike proteins is essential for elucidating the mechanism of packaging and maturation of virions during release from the cellular membrane. We tested our antibodies’ performance in IF using HeLa cells overexpressing trimeric spike protein with the transmembrane domain. Consistent with their performance in the above-mentioned assays (Fig. 2A and 2B), both S1D7 and S3D8 could detect spike proteins expressed homogeneously on the apical side of HeLa cells with a high signal-to-noise ratio (Fig. 2C and S2B). However, their localization pattern is different from a previous report that observed spike proteins exclusively in the Golgi during SARS-CoV-1 infection (Stertz et al., 2007). One possible reason for the difference could be that the spikes were expressed with no other viral proteins (see also Fig. 4B). Mouse hepatitis coronavirus spike protein localizes in the endoplasmic reticulum-Golgi intermediate compartment (ERGIC) in a membrane (M) protein dependent manner. In contrast, when expressed by itself, the spike had a faint reticular appearance (Artika, Dewantari, & Wiyatno, 2020; Opstelten, Raamsman, Wolfs, Horzinek, & Rottier, 1995).

**Figure 3.**
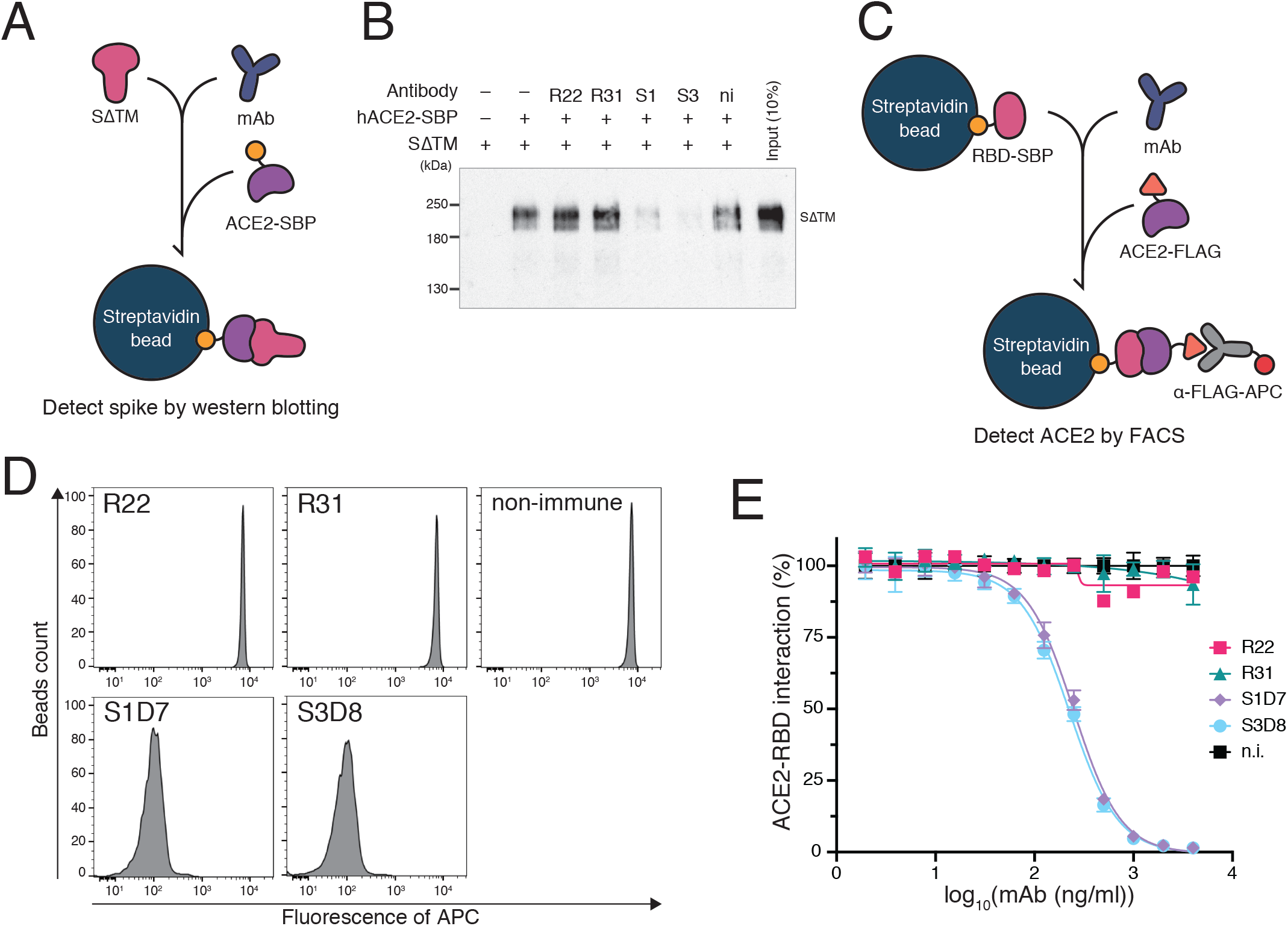
Inhibition of ACE2-spike interaction by S1D7 and S3D8. A. A schematic of the spike pull-down assay designed to evaluate inhibition of ACE2-spike binding by monoclonal antibody. Spike glycoprotein lacking TM domain (SΔTM) was mixed with a monoclonal antibody. ACE2-SBP was applied to capture SΔTM onto streptavidin beads competitively. Captured SΔTM was detected by WB as a measurement of the antibody’s inhibitory ability. S1, S1D7; S3, S3D8; ni, non-immune mouse IgG. B. WB of spike pull-down assay using antibody R52. In the presence of clones S1D7 and S3D8, ACE2 was not able to pull down SΔTM. C. Schematic of bead-based neutralization assay designed to quantify inhibition of ACE2-RBD binding by monoclonal antibody. RBD-SBP glycoprotein immobilized on streptavidin beads was mixed with a monoclonal antibody. ACE2-FLAG was applied to bind competitively with RBD. ACE2-RBD binding was quantified by measuring the signal given by an anti-FLAG antibody conjugated with APC fluorophore using FACS. D. One set of representative FACS results of a bead-based neutralization assay in the presence of 4 µg/mL monoclonal antibodies. Clones S1D7 and S3D8 significantly inhibited ACE2-RBD interaction, shown as lowered fluorescence intensity of APC. E. Binding profiles of potent neutralizing antibodies. ni, non-immune mouse IgG. Error bars indicate standard deviation (n=3). Clones R22 and R31 showed no inhibition of ACE2-RBD binding, while S1D7 and S3D8 inhibited ACE2-RBD interaction at lower ng/mL levels.

**Figure 4.**
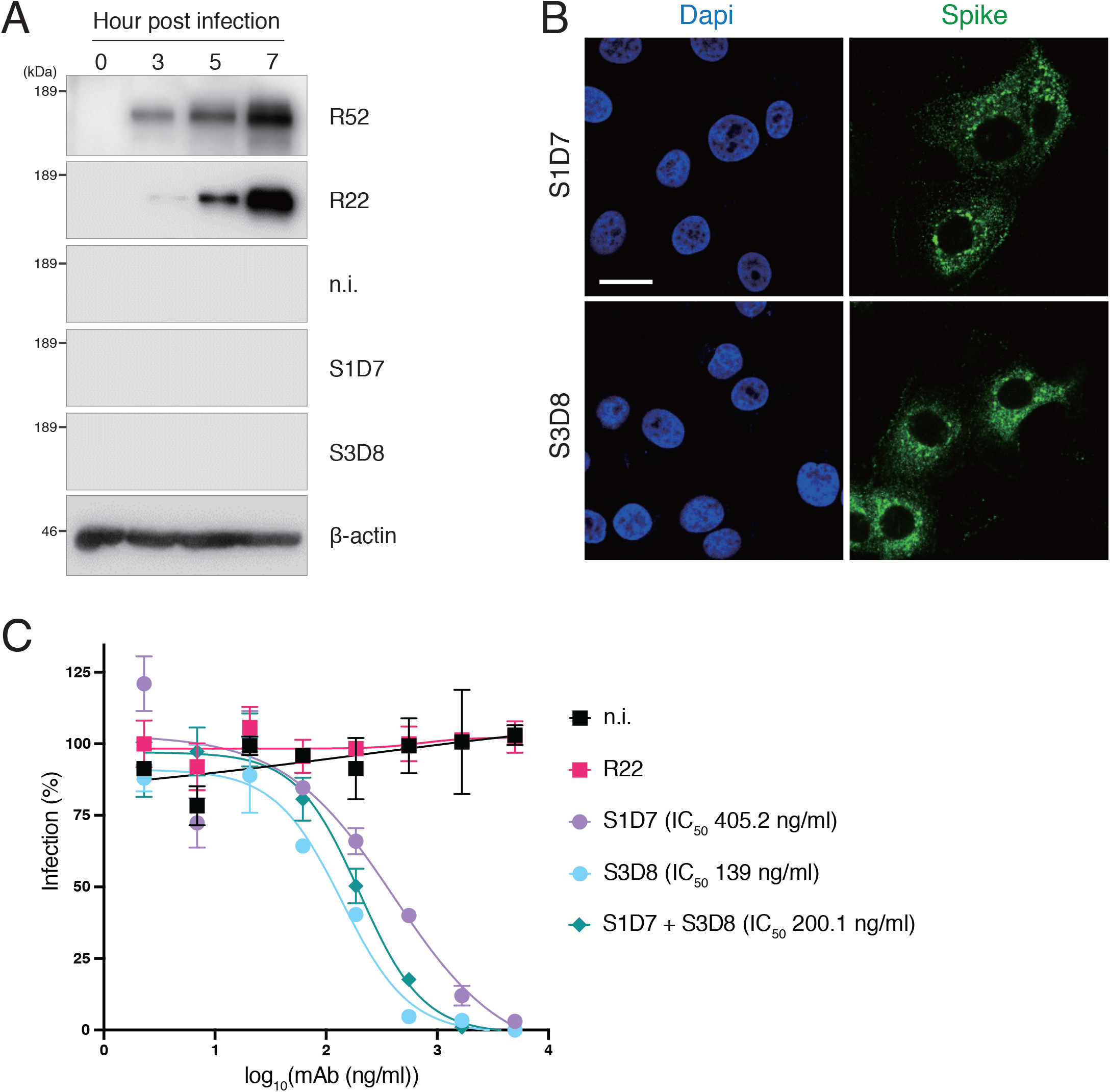
S1D7 and S3D8 neutralized SARS-CoV-2 infection. A. Spike glycoprotein was expressed in VeroE6/TM2 cells during SARS-CoV-2 infection. Spike glycoproteins were detected by western blots using anti-spike antibodies R22 and R52. B. Immunofluorescence staining of spike glycoprotein expressed in VeroE6/TM2 cells infected with SARS-CoV-2 at 7 h post-infection. Scale bar, 20 µm. C. S1D7 and S3D8 are capable of neutralizing live virus infections. Although clone R22 failed to protect VeroE6/TM2 cells from SARS-CoV-2 infection, S1D7 and S3D8 blocked SARS-CoV-2 infection significantly with IC_50_ values of 405.2 ng/mL and 139 ng/mL, respectively. S1D7 and S3D8 cocktail showed intermediate neutralizing activity (200.1 ng/mL). Error bars indicate standard deviation (n=3).

### ACE2-Spike binding inhibition of the monoclonal antibodies

The manner in which antibodies bind and pull down spike glycoproteins in an IP experiment resembles the process of antibody-mediated neutralization, where spike-ACE2 interaction is intercepted by competitive binding between neutralizing antibodies and spike glycoprotein. The performance of our antibodies in IP experiments prompted us to examine whether they were capable of inhibiting spike-ACE2 binding or even neutralizing SARS-CoV-2 infection. First, we performed a spike pull-down assay in which the spike glycoprotein was pulled down by ACE2 in the presence of monoclonal antibodies (Fig. 3A and S3A). Clones S1D7 and S3D8 clearly inhibited spike-ACE2 binding, as shown by the dimmed spike signal in WB (Fig. 3B). To quantify the inhibition ability, we performed a bead-based neutralization assay by measuring the amount of ACE2 bound to RBD beads after blocking with monoclonal antibodies (Fig. 3C). Antibodies R22 and R31 showed no disruption of ACE2-RBD interaction, whereas S1D7 and S3D8 showed robust hindrance of ACE2-RBD binding with IC_50_ values of 248.2 ng/mL and 225.6 ng/mL, respectively (Fig. 3D and 3E). S1D7 and S3D8’s abilities to inhibit spike-ACE2 binding was consistent with their superior performance in IP experiments.

### S1D7 and S3D8 showed neutralizing activity against SARS-CoV-2

Next, we asked whether our antibodies inhibit SARS-CoV-2 infection in VeroE6/TMPRSS2 (TM2) cells, which is susceptible to SARS-CoV-2 infection compared with the parental VeroE6 cell line by expressing TMPRSS2 (Matsuyama et al., 2020). In WB, antibodies R52 and R22, but not S1D7 and S3D8, could detect spike glycoprotein along with the progression of SARS-CoV-2 infection in VeroE6/TM2 cells (Fig. 4A). On the other hand, S1D7 and S3D8 were applicable to IF in infected VeroE6/TM2 cells. Spike showed a punctate distribution pattern in the perinuclear region resembling ER and ERGIC (Sadasivan, Singh, & Sarma, 2017) (Fig. 4B). The subcellular localization of spike resembled that of the N protein in Vero cells infected with SARS-CoV-1 (Stertz et al., 2007), suggesting assembly of SARS-CoV-2 virion in the cytoplasm. We then conducted a live virus neutralization assay to examine whether clones S1D7 and S3D8 inhibit the live virus infection. As expected, although clone R22 failed to protect VeroE6/TM2 from SARS-CoV-2 infection, S1D7 and S3D8 blocked SARS-CoV-2 infection significantly with IC_50_ values of 405.2 ng/mL and 139 ng/mL, respectively, even at relatively high titers of 1500 TCID_50_ (Fig. 4C, Table 1). A cocktail of S1D7 and S3D8 showed intermediate neutralizing activity (200.1 ng/mL), suggesting that S1D7 and S3D8 share an inhibitory mechanism.

**Table 1.** Monoclonal antibodies neutralizing SARS-CoV-2 infection.

| Clone | Host species | Cell line | Virus titer |  |  | IC<sub>50</sub> (ng/ml) | Reference |
| --- | --- | --- | --- | --- | --- | --- | --- |
|  |  |  | TCID<sub>50</sub> | PFU | FFU |  |  |
| BD-368-2 | Human | Vero | - | 100 | - | 15 | Cao *et al.*, Cell |
| 0304-3H3 |  | VeroE6 | 100 | - | - | 110 | Chi *et al.*, Science |
| 1B07* |  | VeroE6 | - | - | 100 | 37 | Hassan *et al.*, Cell |
| P2C-1F11 |  | VeroE6 | - | - | 600 | 15 | Ju *et al.*, Nature |
| S309 |  | VeroE6 | - | - | 100 | 79 | Pinto *et al.*, Nature |
| CC6.33 |  | HeLa-ACE2 | - | 150 | - | 39 | Roger *et al.*, Science |
| CB6 |  | VeroE6 | 100 | - | - | 36 | Shi *et al.*, Nature |
| 47D11 |  | VeroE6 | 500 | - | - | 570 | Wang C *et al.*, Nat Commun |
| B38 |  | Vero | 100 / 200 | - | - | 177 / 1,967 | Wu *et al.*, Science |
| COV2-2196 |  | VeroE6 | - | - | 100 | 15 | Zost *et al.*, Nature |
| S1D7 | Mouse | VeroE6/TM2 | 1500 | - | - | 405.2 | This paper |
| S3D8 |  | VeroE6/TM2 |  |  |  | 139 |  |

\*1B07 is a chimeric monoclonal antibody which combines mouse Fv and human Fc.

## Discussion

Emerging SARS-CoV-2 is a global public health threat to society, which is predicted to be long-term for several years (Kissler, Tedijanto, Goldstein, Grad, & Lipsitch, 2020). Although there are multiple ongoing endeavors to develop neutralizing antibodies, vaccines, and drugs against the virus (Callaway, 2020; Riva et al., 2020), the lack of adequate, licensed countermeasures underscores the need for a more detailed and comprehensive understanding of the molecular mechanisms underlying the pathogenesis of the virus (Artika et al., 2020). Fundamental knowledge has significant implications for developing countermeasures against the virus, including diagnosis, vaccine design, and drug discovery. Due to the above reasons, and our experiences with routine antibody productions (Iwasaki et al., 2016; Murano et al., 2019), we have established and characterized mouse monoclonal antibodies that can be used to dissect the molecular mechanism of the virus life cycle. These antibodies would serve as a reliable toolset for basic research investigating the expression profile and subcellular localization of spike glycoprotein during viral entry, replication, packaging, and budding. These antibodies could help to identify novel host factors interacting with spike glycoprotein when used in IP in combination with mass spectrometry. Therefore, advancement in basic research would accelerate the discovery of drugs targeting virus transmission.

Since passive immunization with neutralizing antibodies has been proposed as a treatment for COVID-19 (Dhama et al., 2020; Jawhara, 2020; S. Jiang et al., 2020; Klasse & Moore, 2020; Ni et al., 2020), research interests have largely focused on cloning human neutralizing antibodies from COVID-19 patients. Our antibodies, S1D7 and S3D8, have been shown to attenuate the interaction of spike proteins with ACE2 and neutralize infection of VeroE6/TM2 cells by SARS-CoV-2. It is worth noting that although their neutralizing activities (IC_50_ of 405.2 ng/ml and 139 ng/ml) appeared to be lower than those of human antibodies reported previously (Fig. 4C and Table 1), the stringency of experimental conditions (relatively high virus titer of 1500 TCID_50_) tend to underestimate neutralizing activities of our antibodies compared to other research groups. Specifically, we used a high multiplicity of live SARS-CoV-2 virus to infect VeroE6/TM2 cells, which are more prone to virus infection than the commonly adopted VeroE6 cell line. Therefore, it is difficult to compare antibody efficacy among them (Tse, Meganck, Graham, & Baric, 2020). In addition to *in vitro* infection, their neutralizing activity *in vivo* should be examined in animal models that recapitulate SARS-CoV-2 disease. Our mouse antibodies will not be applicable for use in clinical treatment, if not chimeric and humanized, due to their immunogenicity (Hansel et al., 2010; Reichert et al., 2005). On the other hand, they may be valuable for investigating the mechanism of immune responses to the virus during passive immunization using mouse models for SARS-CoV-2 infection (Bao et al., 2020; Dinnon et al., 2020; Hassan et al., 2020; Israelow et al., 2020; R. D. Jiang et al., 2020; Winkler et al., 2020). They could show stable performance due to lot-to-lot consistency and act as benchmarks for other antibodies and drug developments.

## Acknowledgements

We thank Ayako Ishida, Mie Kobayashi, and Yasuyuki Kurihara for technical assistance and advice on the production of antibodies. This work was supported by the Keio University Global Research Institute (KGRI) COVID-19 Pandemic Crisis Research Grant (to M.T., H.S., and K.M.) and the Keio Donner Project, which is devoted to Shibasaburo Kitasato, the founder of Keio University School of Medicine. The project was built under the leadership of the top officials of Keio University, Tsutomu Takeuchi (Vice-President), Masayuki Amagai (Dean of the School of Medicine), Yuko Kitagawa (Hospital Director General), and Hideyuki Saya (Project Coordinator). This study was also supported in part by AMED (JP18fk0108076 to A.K.), NOMURA Microbial Community Control Project in ERATO of the Japan Science and Technology Agency (to A.K.), and Research Support Program to Tackle COVID-19 Related Emergency Problems, University of Tsukuba (to A.K.).

## Author contributions

Y.G., H.S., and K.M. conceived the project, designed the experiments and wrote the manuscript. Y.G., A.K., M.T., and K.M. performed the experiments. All authors analyzed the data and contributed to the preparation of the manuscript.

## Competing interests

The authors declare no competing interests.

## Materials and Methods

### Expression and purification of proteins in human cells

Synthetic DNA sequences encoding SARS-CoV-2 spike protein ectodomain (SΔTM, residue 1-1208; strain Wuhan-hu-1; GenBank: QHD43416.1) and RBD (residue 319-591; strain Wuhan-hu-1; GenBank: QHD43416.1) fused with an N-terminal signal peptide, a C-terminal trimerization motif, an HRV3C cleavage site, an SBP purification tag, and an 8xHis-tag were inserted into pEFx mammalian expression vector. S1/S2 (682-RRAR-685) and S2’ (986-KV-987) cleavage sites of spike protein were mutated (682-GSAS-685, 986-PP-987, respectively) to prevent protease cleavage. The codon composition of DNA fragments was optimized and synthesized for protein expression in human cells (FASMAC). Full-length spike, human ACE2-SBP (residue 1-708; NP_001358344), and ACE2-FLAG were synthesized and cloned into pcDNA3.4 vector (Thermo Fisher). Recombinant proteins were prepared by Expi293 Expression System (Thermo Fisher) according to the manufacturer’s instruction. They were secreted into culture medium supernatant of the Expi293F cells, and then affinity purified by Streptavidin Sepharose High Performance (Cytiva). Purification tags were removed by treating recombinant proteins with HRV3C protease (TaKaRa) and cOmplete™ His-tag Purification Resin (Roche). Purity and glycosylation of recombinant proteins were examined by PNGase F (N-Zyme Scientifics) treatment followed by SDS-PAGE and Coomassie staining. The containment measures for the living modified organisms in all experiments were confirmed by the Ministry of Education, Culture, Sports, Science and Technology of Japan on April 27, 2020.

### Expression and purification of proteins in *E. coli*

The DNA sequence encoding SARS-CoV-2 spike protein RBD (residue 410-580; strain Wuhan-hu-1; GenBank: QHD43416.1) was amplified from a nasopharyngeal swab of a patient treated in the Keio University Hospital, and in-framed inserted into pGEX-5X-1 and pMAL-c2G *E. coli* expression plasmid, downstream of GST-tag and MBP-tag encoding sequence respectively. Sample collection is approved by Keio University Bioethics Committee with the number 20200063. Recombinant proteins were expressed in overnight 16°C cultured BL21(DE3)pLysS competent cells transformed by corresponding vector under induction of 1 mM Isopropyl β-D-1-thiogalactopyranoside (IPTG). MBP-tagged RBD was affinity purified by Amylose Resin (NEB) according to manufacturer’s instructions; GST-tagged RBD was affinity purified by Glutatione Sepharose 4B (Cytiva) according to manufacturer’s instructions. Purity of purified recombinant proteins were examined by SDS-PAGE followed by Coomassie staining.

### Cell cultures

The mouse myeloma cell line SP2/0-Ag14 (RCB0209) was provided by the Riken Bioresouces Center (Tsukuba, Japan). The cells were cultured in RPMI 1640 (Nissui) supplemented with 10% heat-inactivated calf serum (Biowest) and 1 ng/mL recombinant human interleukin 6 (IL-6, PeproTech). HeLa and 293T cells were cultured in DMEM (Nacalai tesque) with 10% fetal bovine serum (Biowest). We maintained hybridoma clones against spike glycoproteins in Hybridoma Serum-free Medium (FUJIFILM Wako) supplemented with 1 ng/mL IL-6.

### Production of monoclonal antibodies

BALB/c mice were immunized twice in 3-week intervals, with the second immunization serving as a booster. Mice were injected intraperitoneally with 100 µL chyle containing 10-50 µg antigen prepared with TiterMax Gold adjuvant (Sigma-Aldrich) according to the manufacturer’s instructions. Four days after boosting, splenocytes of immunized mice were collected by grinding the spleens in RPMI 1640 medium. Splenocytes (1×10^8^) were immediately mixed with 5×10^7^ SP2/0 myeloma cells and fused using an electro cell fusion generator ECFG21 (NepaGene) according to the manufacturer’s instructions. After fusion, cells were cultured in HAT medium (RPMI 1640 supplemented with 10% calf serum containing HT supplement (Gibco) and 0.4 µM aminopterin (Sigma-Aldrich)) for 10 days to select hybridomas. Hybridomas were subsequently screened by ELISA, in which RBD glycoproteins were generated from the Expi293F expression system. We performed western blotting and immunoprecipitation for further screening and subjected to monoclonization by serial dilution. For antibody production, monoclonal hybridomas were cultured in Hybridoma Serum-Free Medium (FUJIFILM Wako) supplemented with IL-6. Monoclonal antibodies were purified from hybridoma culture supernatants using Thiophilic-Superflow Resin (Clontech) or Ab-Capcher MAG2 (ProteNova) according to the manufacturer’s instructions. The isotype of antibodies was determined using the IsoStrip Mouse Monoclonal Antibody Isotyping Kit (Roche).

### Western blotting and immunoprecipitation

SΔTM and RBD glycoproteins were resolved on SDS-PAGE and transferred onto a nitrocellulose membrane (Amersham Protran, GE Healthcare). Lysates of 293T cells transfected with plasmids encoding full-length spike glycoproteins was also separated by SDS-PAGE for WB. The membrane was blocked in 1% nonfat skim milk and then incubated in 1 µg/mL anti-spike antibodies for 1h at room temperature. After three times washing in PBS-T (0. 1% Tween-20), the membrane was incubated in 1:5000 dilution of the peroxidase-conjugated sheep anti-mouse IgG secondary antibody (MP Biomedicals) for 30 min at room temperature. Signals were detected using ECL Western Blotting Detection Reagents (GE Healthcare).

For immunoprecipitation assay, 1 µg of purified antibodies was conjugated to 10 µl Dynabeads Protein G (Thermo Fisher) for 30 min at room temperature, followed by washing twice in IP buffer (20 mM Tris-HCl(pH 7.4), 150 mM NaCl, 0.1% NP-40). Antibody conjugated beads were incubated with 100 ng SΔTM in 50 µl IP buffer for 2 hours at room temperature. Beads were washed three times in IP buffer and eluted with SDS-PAGE loading dye at 95°C for 5 min. Immunoprecipitation of SΔTM was examined by SDS-PAGE followed by western blotting using antibody R52.

### Immunofluorescence

Before performing immunofluorescence, HeLa cells seeded on cover glasses were transfected with plasmids encoding full length SARS-CoV-2 spike protein for 2 days using Lipofectamine 2000 (Thermo Fisher). Cells were fixed with 2% formaldehyde in PBS for 10 min at room temperature, washed in PBS-T once, and permeabilized with 0.1% Triton X-100 in PBS for 10 min at room temperature. Cells were blocked by 1% non-fat skim milk in PBS-T for 10 min, then incubated with 0.5 µg/mL antibody for 1 h at room temperature. After three times wash in PBS-T, cells were incubated in 1:500 diluted Alexa Fluor 488 conjugated goat anti-mouse IgG secondary antibody (Thermo Fisher) and 1 µg/mL DAPI solution for 30 min at room temperature. The cover glasses were mounted with Prolong Glass Antifade Mountant (Thermo Fisher) overnight at room temperature before observing. The fluorescence images were taken with Keyence BZ-X810 fluorescence microscope and Olympus FV3000 confocal laser scanning microscope.

### ELISA of antibody binding to SARS-CoV-2 spike protein

Nunc MaxiSorp™ flat-bottom 96-well plates (Thermo Fisher) were coated with 170 ng SΔTM in 50 µl PBS overnight at 4°C, then blocked at room temperature for 1 hour by applying 200 µl of 3.75% BSA in PBS-T. Monoclonal antibodies starting from 100 µg/mL were four-folds serial diluted with blocking buffer to 12 gradients and incubated with plates for 1 hour at room temperature, followed by incubation with horseradish peroxidase (HRP) conjugated sheep anti-mouse secondary antibody (MP Biomedicals) 1:5000 diluted in blocking buffer for 30 min at room temperature. Plates were incubated for 15 min at room temperature with 1-Step Turbo TMB-ELISA Substrate Solution (Thermo Fisher), then terminated with equal volume of 1M phosphoric acid. Signal was quantified by measuring absorbance at 450 nanometer using iMark Microplate Absorbance Reader (Bio-Rad Laboratories). Half-maximum effective concentration (EC_50_) was calculated by non-linear regression analysis of absorbance curve.

### ACE2-binding inhibition assay

For spike pull-down assay, SΔTM glycoprotein was incubated with 1 µg anti-spike antibody in 50 µl binding buffer (PBS supplemented with 0.1% NP-40) at room temperature for 1 hour, then 3 µg of ACE2-SBP recombinant protein was applied the reaction for 1 hour. The ACE2-SBP was pull-down by 10µl Dynabeads M-270 Streptavidin (Thermo Fisher) for 30 min at room temperature, followed by washing twice with binding buffer and elution with SDS-PAGE loading dye at 95°C for 5 min. ACE2-Spike binding inhibition was examined by SDS-PAGE, followed by WB using antibody R52.

For bead-based neutralization assay, 20 µl of Streptavidin beads were incubated with 4 µg of RBD-SBP in 100 µl of TBSTx (TBS supplemented with 1% TritonX-100) overnight at 4°C with shaking. After washing, beads were incubated with diluted antibodies for 20 min at 4°C, washed, incubated with 4 µg/mL of ACE2-FLAG for 20 min at 4°C, washed, and incubated with an anti-DYKDDDDK antibody conjugated with APC fluorophore (MBL) for 20 min at 4°C. After the final wash, the mean fluorescence intensity (MFI) of beads was analyzed by a FACS Verse (BD). The relative MFI of beads was calculated by normalization using the MFI of beads incubated with non-immune mouse IgG.

### Virus neutralization assay

SARS-CoV-2 virus (obtained from the National Institute of Infectious Diseases) was prepared from culture fluids harvested from infected VeroE6/TMPRSS2 cells (JCRB Cell Bank, JCRB1819) (Matsuyama et al., 2020). The virus titer was 3 × 10^7^ TCID_50_/mL. The virus solution containing 1500 TCID_50_ was incubated with each antibody at concentrations of serial threefold dilutions starting from 5 µg/mL. After incubating at room temperature for 1 h, the antibody-treated virus solution was mixed with VeroE6/TMPRSS2 cells in glass-bottom 96-well plates. At 7 h post-infection, cells were fixed in 4% PFA and subjected to indirect immunofluorescence assays using S1D7 antibody as described above. The number of infected cells were imaged and analyzed using ArrayScan (Thermo Fisher). Mouse anti-FLAG M2 antibody (Sigma) was also used as a control. Experiments with SARS-CoV-2 were performed in a biosafety level 3 (BSL3) containment laboratory at University of Tsukuba.

#### Supplemental Figure Legends

**Figure S1.**
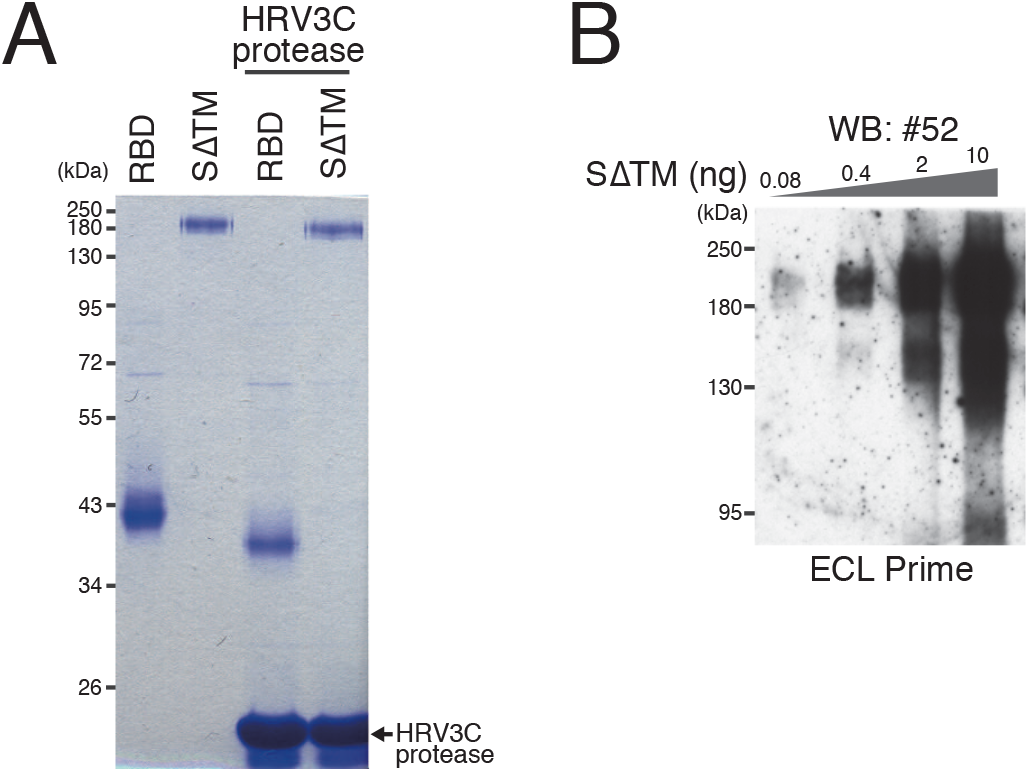
A. Recombinant spike glycoproteins were treated with HRV3C protease to remove SBP-tag before immunizing mice. B. Clone R52 showed the highest performance on western blotting among our antibodies and detected even 0.08 ng SΔTM glycoprotein.

**Figure S2.**
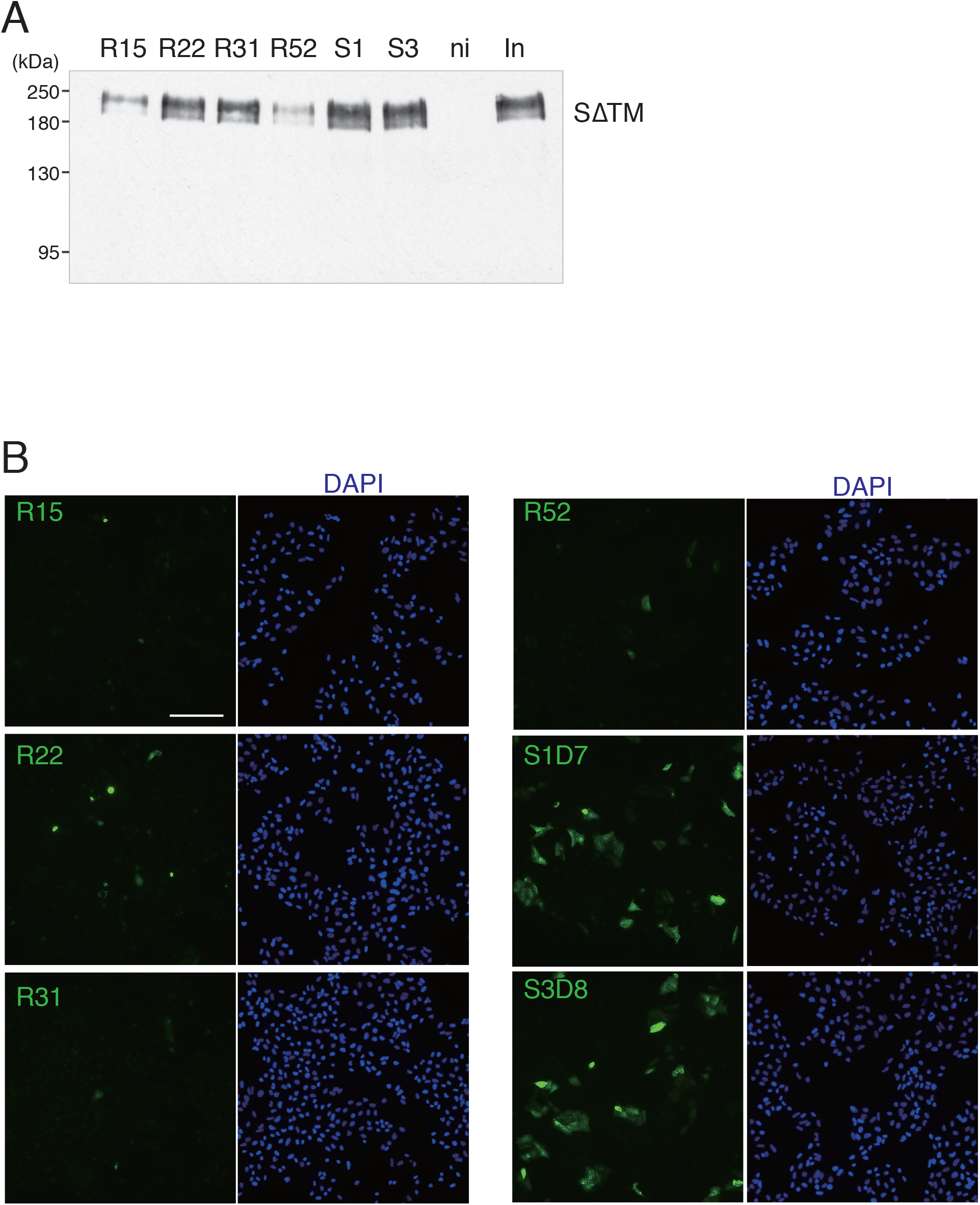
A. Monoclonal antibody clones S1D7 and S3D8 maintain high efficiency even in the presence of 0.1% SDS. S1, S1D7; S3, S3D8; ni, non-immune IgG; In, input. B. Immunofluorescence (IF) staining of spike glycoprotein expressed in HeLa cells with all six monoclonal antibodies. S1D7 and S3D8 showed higher performance in IF. Images were captured using a Keyence BZ-X810 fluorescence microscope. Scale bar, 200 µm.

**Figure S3.**
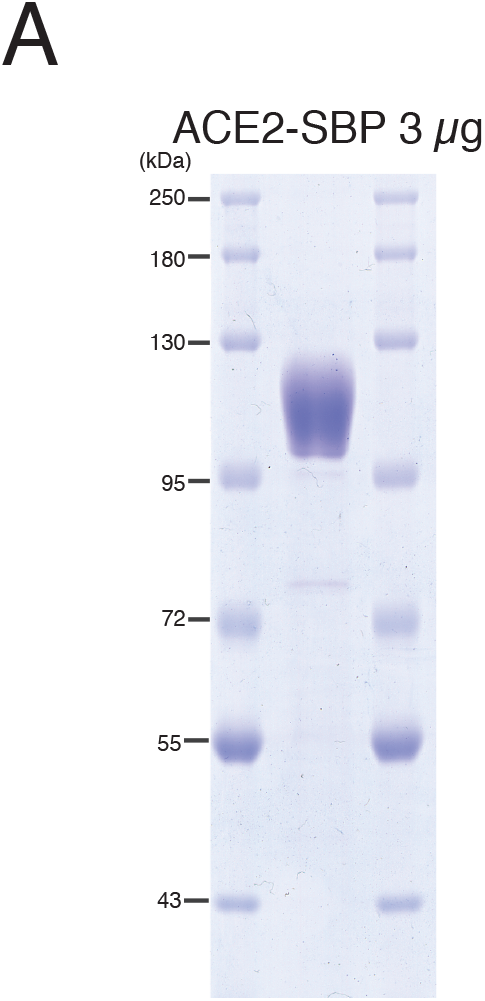
A. ACE2-SBP protein was purified from the culture supernatant of Expi293F cells transfected with a plasmid encoding ACE2-SBP.

## Notes

### Competing Interest Statement

The authors have declared no competing interest.

